# Geographic heterogeneity impacts drug resistance predictions in *Mycobacterium tuberculosis*

**DOI:** 10.1101/2020.09.17.301226

**Authors:** Guo Liang Gan, Matthew H. Nguyen, Elijah Willie, Mohammad H. Rezaie, Brian Lee, Cedric Chauve, Maxwell Libbrecht, Leonid Chindelevitch

## Abstract

The efficacy of antibiotic drug treatments in tuberculosis (TB) is significantly threatened by the development of drug resistance. There is a need for a robust diagnostic system that can accurately predict drug resistance in patients. In recent years, researchers have been taking advantage of whole-genome sequencing (WGS) data to infer antibiotic resistance. In this work we investigate the power of machine learning tools in inferring drug resistance from WGS data on three distinct datasets differing in their geographical diversity.

We analyzed data from the Relational Sequencing TB Data Platform, which comprises global isolates from 32 different countries, the PATRIC database, containing isolates contributed by researchers around the world, and isolates collected by the British Columbia Centre for Disease Control in Canada. We predicted drug resistance to the first-line drugs: isoniazid, rifampicin, ethambutol, pyrazinamide, and streptomycin. We focused on the genes which previous evidence suggests are involved in drug resistance in TB.

We called single-nucleotide polymorphisms using the Snippy pipeline, then applied different machine learning models. Following best practices, we chose the best parameters for each model via cross-validation on the training set and evaluated the performance via the sensitivity-specificity tradeoffs on the testing set.

To the best of our knowledge, our study is the first to predict antibiotic resistance in TB across multiple datasets. We obtained a performance comparable to that seen in previous studies, but observed that performance may be negatively affected when training on one dataset and testing on another, suggesting the importance of geographical heterogeneity in drug resistance predictions. In addition, we investigated the importance of each gene within each model, and recapitulated some previously known biology of drug resistance. This study paves the way for further investigations, with the ultimate goal of creating an accurate, interpretable and globally generalizable model for predicting drug resistance in TB.

**Author summary:** Drug resistance in pathogenic bacteria such as *Mycobacterium tuberculosis* can be predicted by an application of machine learning models to next-generation sequencing data. The received wisdom is that following standard protocols for training commonly used machine learning models should produce accurate drug resistance predictions.

In this paper, we propose an important caveat to this idea. Specifically, we show that considering geographical diversity is critical for making accurate predictions, and that different geographic regions may have disparate drug resistance mechanisms that are predominant. By comparing the results within and across a regional dataset and two international datasets, we show that model performance may vary dramatically between settings.

In addition, we propose a new method for extracting the most important variants responsible for predicting resistance to each first-line drug, and show that it is to recapitulate a large amount of what is known about the biology of drug resistance in *Mycobacterium tuberculosis*.

## Introduction

Tuberculosis (TB) is a serious infectious disease caused by *Mycobacterium tuberculosis* (MTB) and one of the 10 leading causes of death in the world, with roughly 1.4 million people dying from the disease every year [1]. Based on the latest WHO TB report, TB incidence is decreasing at approximately 2% each year, and about 16% of the cases of the disease result in death. More efforts are needed to reach the targets of at most a 5% case fatality ratio and at most a 10% TB treatment failure rate adopted by the End TB Strategy [2]. In order to achieve these milestones, a lot of research has focused on addressing one of the most pressing issues in TB, drug resistance. MTB is susceptible to acquiring resistance, which poses challenges for an effective treatment of the disease. 600,000 new cases of resistance to rifampicin, one of the key first-line drugs, were reported in 2016, and 490,000 of these cases were multi-drug resistant, defined as resistance to rifampicin and isoniazid, the two most effective first line-drugs [1]. Drug resistance arises and continues to spread largely due to two reasons: improper TB drug treatment, whereby poor quality drugs, premature treatment discontinuation, and inappropriate drug regimens apply selective pressure on the MTB bacteria within patients that leads to drug resistance, and person-to-person transmission, whereby these drug-resistant MTB strains are spread to new patients [1].

The gold standard in detecting drug resistance is phenotypic drug susceptibility testing via bacterial culture, whereby MTB is cultured in a Lowenstein-Jensen medium and exposed to a particular drug, and its response to the drug is then observed. This process is slow and can require up to two months before results are obtained. A faster, also commonly used tool for drug resistance detection is commercial genotypic line probe assays such as XPert MTB/RIF and GenoType MTBDRplus. However, such assays only predict resistance to isoniazid and/or rifampicin; they also only consider common mutations associated with resistance, and therefore have lower sensitivity. Due to recent advancements in whole-genome sequencing (WGS) technology, researchers are able to sequence the whole genome of bacteria within a few hours at a much lower cost while extracting more information about genotypic resistance markers. Mutation libraries and databases based on single-nucleotide polymorphisms (SNPs) have been constructed to detect drug resistance; these are included in tools such as TBProfiler [3], TBDreaMDB [4], PhyResSE [5] and Mykrobe [6]. However, these libraries are not able to capture the interactions between different resistance markers in predicting resistance; moreover, they are highly dependent on the diversity of the sequenced samples.

Machine learning and deep learning have been successfully applied to different areas of research such as natural language processing, computer vision, and more recently bioinformatics, including GWAS and cancer prognosis prediction [7, 8]. Leveraging the abundance of WGS data, researchers have also applied machine learning tools to predicting antibiotic resistance in MTB; for instance, in [9] they applied machine learning models such as logistic regression, support vector machines, and naive Bayes classifiers to predict resistance to 8 drugs from WGS data; in [10], the authors presented a deep learning model, a multi-task wide and deep neural network for drug resistance prediction, where they combined logistic regression and various deep learning techniques. Both of these approaches have demonstrated promising results relative to curated libraries in terms of specificity and sensitivity.

Previous work has focused exclusively on the prediction aspect and has not fully explored the insights which could be obtained from the trained machine learning models themselves, namely, a feature importance analysis. Such an analysis is especially important for biological problems, where practitioners such as clinicians or public health professionals may be interested in studying the interpretable factors, such as the genes or SNPs, which are involved in determining resistance. Moreover, previous work has not directly demonstrated the robustness of these models in making predictions on datasets of different diversity.

In this work, we investigate the robustness of machine learning models for MTB drug resistance prediction by training them on multiple datasets of different diversity and composition. We demonstrate the pitfalls with regards to the generalizability of models trained on homogeneous datasets with low diversity and regional data. Moreover, we investigate more in depth those genes that are highly associated with determining resistance by using different methods to determine which genes are of particular importance for each model in determining drug resistance.

### Datasets

In this work, we focus on three geographically distinct datasets: the Relational Sequencing TB Data Platform (ReSeqTB) [11], the Pathosystems Resource Integration Center (PATRIC) database [12], and a collection of isolates from the British Columbia Centre for Disease Control (BCCDC). ReSeqTB is a consortium dedicated to the development of next-generation TB diagnostics, and the dataset comprises global isolates from 32 different countries. The PATRIC database contains isolates from around the world. We have removed the BCCDC isolates from ReSeqTB. There is no overlap between ReSeqTB and PATRIC. The BCCDC dataset is considered to be homogeneous, since it originates from only one location, while the ReSeqTB and PATRIC datsets are heterogeneous. Since all three datasets contain phenotypes for five first-line drugs - isoniazid (INH), rifampicin (RIF), ethambutol (EMB), pyrazinamide (PZA) and streptomycin (SM) - we focus our analysis on those. In order to minimize cross-dataset variation, we only downloaded raw WGS sequences of each isolate from NCBI’s Sequence Read Archive and called variants ourselves instead of using the provided variants. We could not control the sequencing technology used for the different datasets (BCCDC are HiSeq sequences, whereas ReSeqTB and PATRIC are a mix of HiSeq, MiSeq, and other sequencing technologies). This contributes to the heterogeneity of the dataset, although we believe these difference are not as significant as the genomic diversity within and between datasets.

For our cross-regional experiments, we used the full genotypic data available from ReSeqTB. The data from three regions with sufficient representation are then analyzed: North America (comprising 306 Canadian and 453 USA samples), Asia (comprising 144 Chinese and 101 Vietnamese samples), and Africa (comprising 436 South African samples, 48 Ugandan samples and 73 Sierra Leone samples).

In this work, we predict drug resistance to single drugs and focus on SNPs within two lists of candidate genes, one with 23 genes and one with 179 genes, both of which were previously reported to be associated with drug resistance in MTB [13, 14]. Both lists of genes can be found in the supporting information. The list of 23 genes is a subset of the list of 179 genes. The set of 23 genes was reported to be associated to the first line and second line drugs, whereas the set of 179 genes is an extension of the former set to other antibiotic drugs. Machine learning algorithms are able to capture intricate relationships between different features and we would like to investiagte the effect on drug resistance prediction when we include more SNPs information from a larger set of genes. The number of samples and SNPs in each dataset is presented in 1. Figures 1 and 2 refer to the number of SNPs used for resistance prediction for different drugs as well as the number of susceptible and resistant samples for each drug. As shown in those figures, the BCCDC and the ReSeqTB datasets are very imbalanced for all the first-line drugs (with more susceptible than resistant cases), and there are 3 to 4 times as many SNPs in the list of 179 genes compared to the list of 23 genes. For PATRIC, there are drugs (INH, PZA, RIF) where the imbalance is in the other direction with a lot more resistant than susceptible cases. In addition, among the first-line drugs we considered, PATRIC and ReSeqTB have many co-occurrences of resistance, with most resistant samples being resistant to 2 or more drugs. The BCCDC dataset, however, has most of its samples resistant to only one drug. The co-occurrence of drug resistance is not necessarily directly related to genetics, however; instead, it is a reflection of the commonly used drug cocktails in the treatment of TB [15]. We weighted the training examples such that the two classes are balanced prior to training (i.e. minority class is weighed more heavily, while majority class is weighed more lightly).

**Fig 1.**
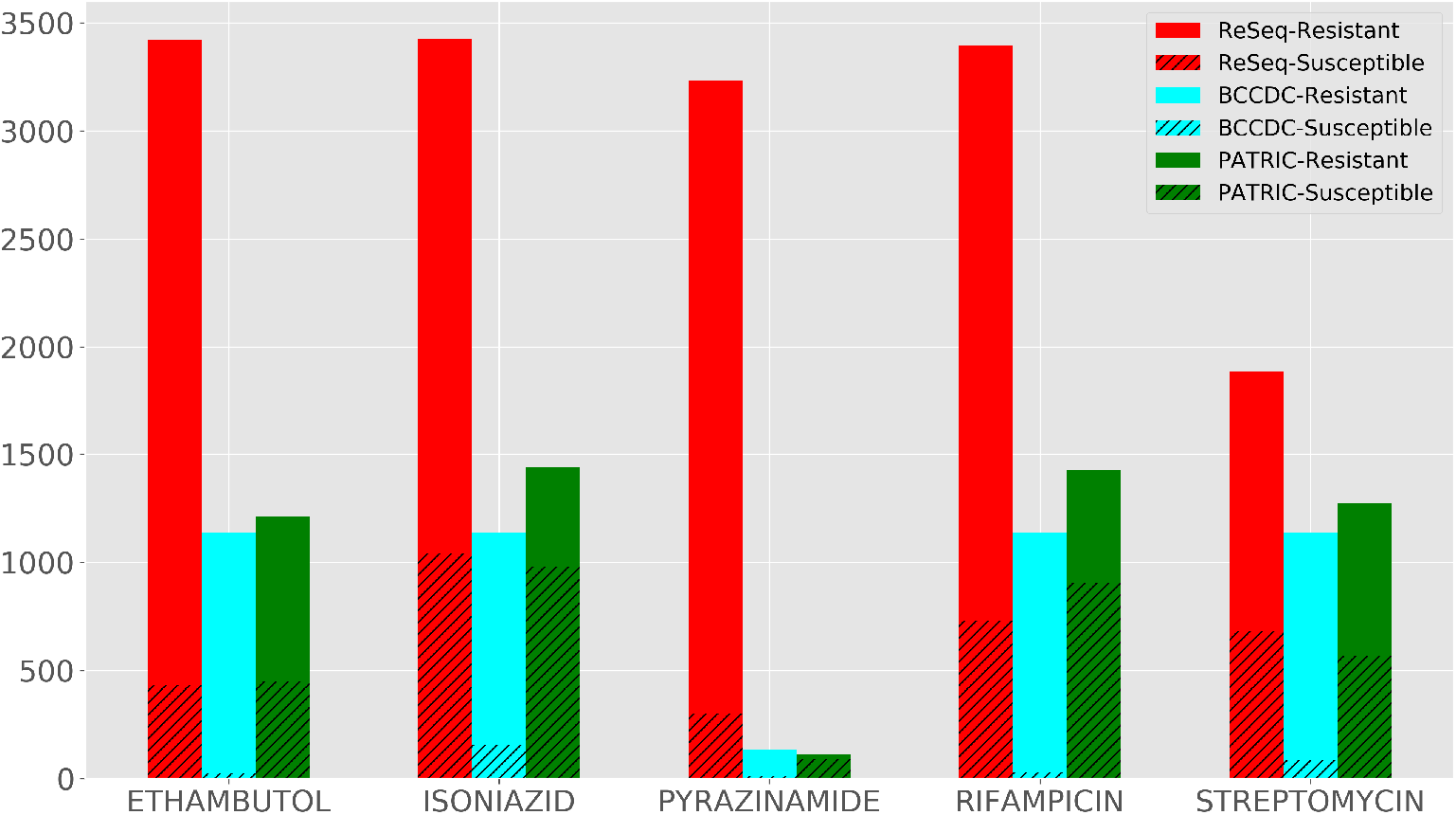
Number of susceptible and resistant samples for different drugs.

**Fig 2.**
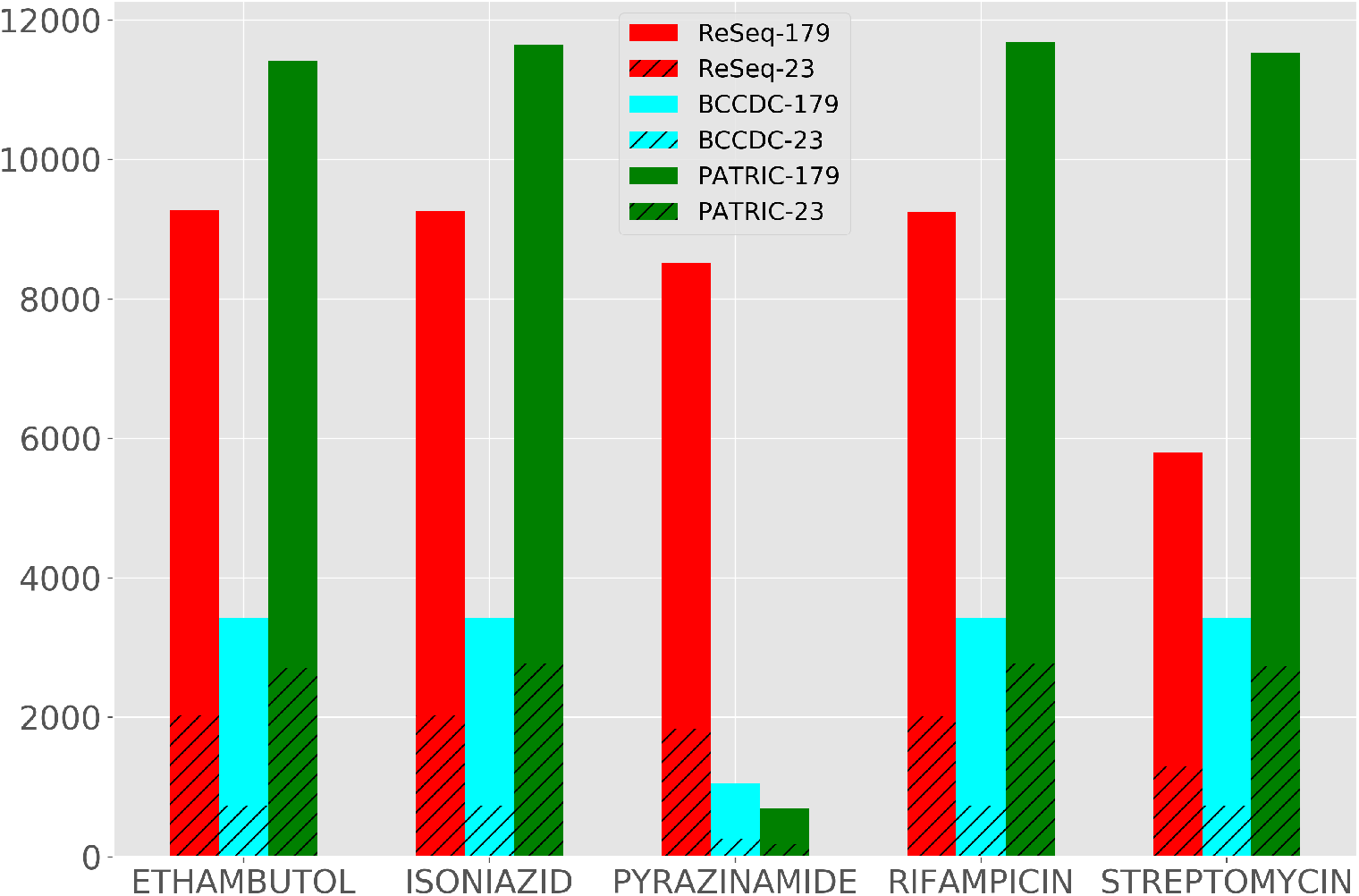
Number of SNPs used to predict resistance to different drugs.

## Methods

Figure 3 illustrates the workflow of our machine learning approach.

**Fig 3.**
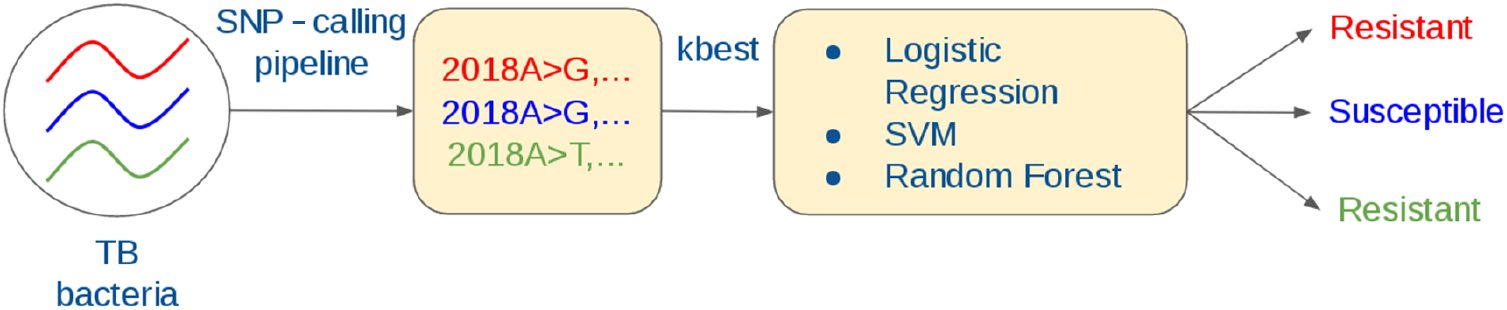
Illustration of the machine learning approach.

**Fig 4.**
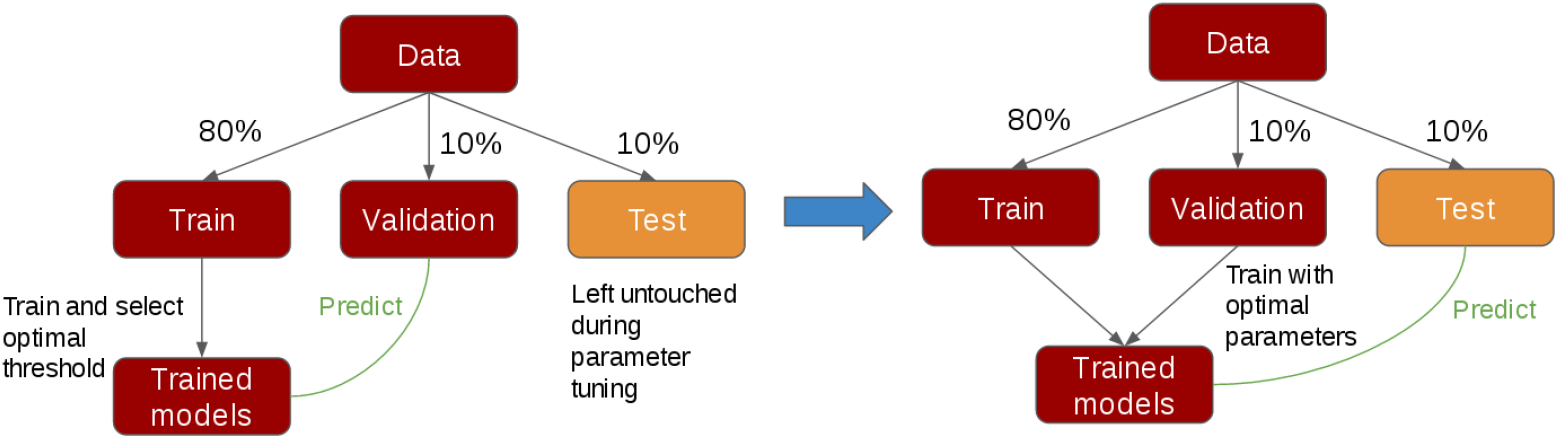
Description of each fold for a 10-fold cross validation process. (Left) The workflow for choosing the best parameters for each model. Models are trained on the training set and an optimal probability threshold chosen for classification. The validation set is used for choosing the best parameters and the testing set was left untouched. (Right) Once the best parameters are found, we train on both the training and validation sets, predict on the testing sets, and report the performance on the testing set for each fold.

### Calling single nucleotide polymorphisms (SNPs)

**For all three datasets, SNPs were called using Snippy** [16], with *Mycobacterium tuberculosis H37Rv* used as the reference genome. We translated the SNPs and the phenotypic data into a feature matrix *X* and a list of label vectors *y_D_* indexed by the drug *D*, respectively. *X* is a binary matrix, where *x_ij_* = 0 indicates that sample *i* does not have the *j*-th SNP, while *x_ij_* = 1 indicates that it does. *y_D_* is a binary vector with 0 indicating susceptibilty and 1 indicating resistance to the given drug D.

For the cross-regional experiments, we used the fully available ReSeqTB genotype data with calls using the UVP pipeline [17]. This was done to ensure more data was available than just the samples committed to a public repository and for which the corresponding accession number was available. This allowed us to obtain SNPs from a total of 6495 isolates through ReSeqTB.

### Cross-validation protocol

#### We performed a 10-fold cross-validation to choose hyperparameters

In order to choose the best set of parameters for each model, we partitioned each fold within the dataset into a training set (80%), a validation set (10%) and a testing set (10%). Only the training and validation sets were used in the process of parameter tuning. We trained the model on the training set and searched for an optimal probability threshold which yielded the highest accuracy on the training set. We measured performance using the area under the Receiver Operating Characteristic curve (AUC). We picked the best parameters for the model based on the average AUC across the 10 folds on the validation set. Once the set of best parameters was found, we retrained the model on the training and validation sets, then predicted on the test set and reported the AUC of the test set for each fold.

### Machine learning classifiers

**We considered the main models used in [9]**, namely, Logistic Regression with *L*_1_ and *L*_2_ penalty (LR_L1 and LR_L2), support vector machine with linear kernel and *L*_2_ penalty (SVM_L2), support vector machine with radial basis kernel (SVM_RBF) and random forests (RF). In our work we additionally investigated the ElasticNet (eNet) model. eNet refers to a logistic regression model which combines *L*_1_ and *L*_2_ penalties.

### Resistance-associated genes

**We investigated resistance-associated genes for each drug based on the machine learning models** using the following two methods:

- *Submatrix permutation test*: For each gene *g* in the list of (23 or 179) candidate genes, we randomly permuted the rows of the submatrix of the input feature matrix *X* whose columns were SNPs within *g*, while leaving the rest of X unchanged. Denoting this permuted matrix as *X_g_*, we then input *X_g_* to the machine learning models for retraining. Let *AUC*(*X*) and *AUC*(*X_g_*) denote the test AUC using *X* and *Xg* as the feature matrix, respectively. We computed Δ_*g*_ = *AUC*(*X*) − *AUC*(*X_g_*). If Δ_*g*_ is positive and large in magnitude, we may conclude that gene *g* is relevant to resistance determination for the drug of interest.
- *Gene-wise partial least square (PLS) regression*: PLS is a regression model which decomposes *X* into a set of components explaining as much as possible of the covariance between X and *y* [18]. We performed PLS on X gene-by-gene and kept only 1 component for each gene. Hence, the PLS-transformed matrix *PLS*(*X*) is an *n × m* matrix, where *n* is the number of samples and *m* is the number of genes considered. We then trained and predicted resistance using the LASSO (LR_L1) [19] and the random forest (RF) models with *X′* := *PLS*(*X*) as the input matrix. By investigating the resulting models, we are able to estimate the importance of each gene for predicting resistance to a particular drug.

## Results

### PCA suggests lineage is not a confounder of drug resistance

We performed Principal Component Analysis (PCA) on the feature matrices for the BCCDC and ReSeqTB datasets to investigate the relations between lineages ^1^ and resistance to isoniazid and pyrazinamide (used as examples). Figure 5 and Figure 6 show the plots of the first 2 PCA components for both datasets respectively, with samples grouped by lineage. For ReSeqTB, the samples can be grouped into clusters reasonably well, one cluster per lineage. Within each cluster, the resistant and susceptible samples are non-separable, which suggests that lineage is not a confounder of resistance. For BCCDC, a similar conclusion applies. No PCA was performed for PATRIC since the associated metadata did not contain lineage information. Similar observations for both datasets suggested that lineage is not a confounder of drug resistance.

**Fig 5.**
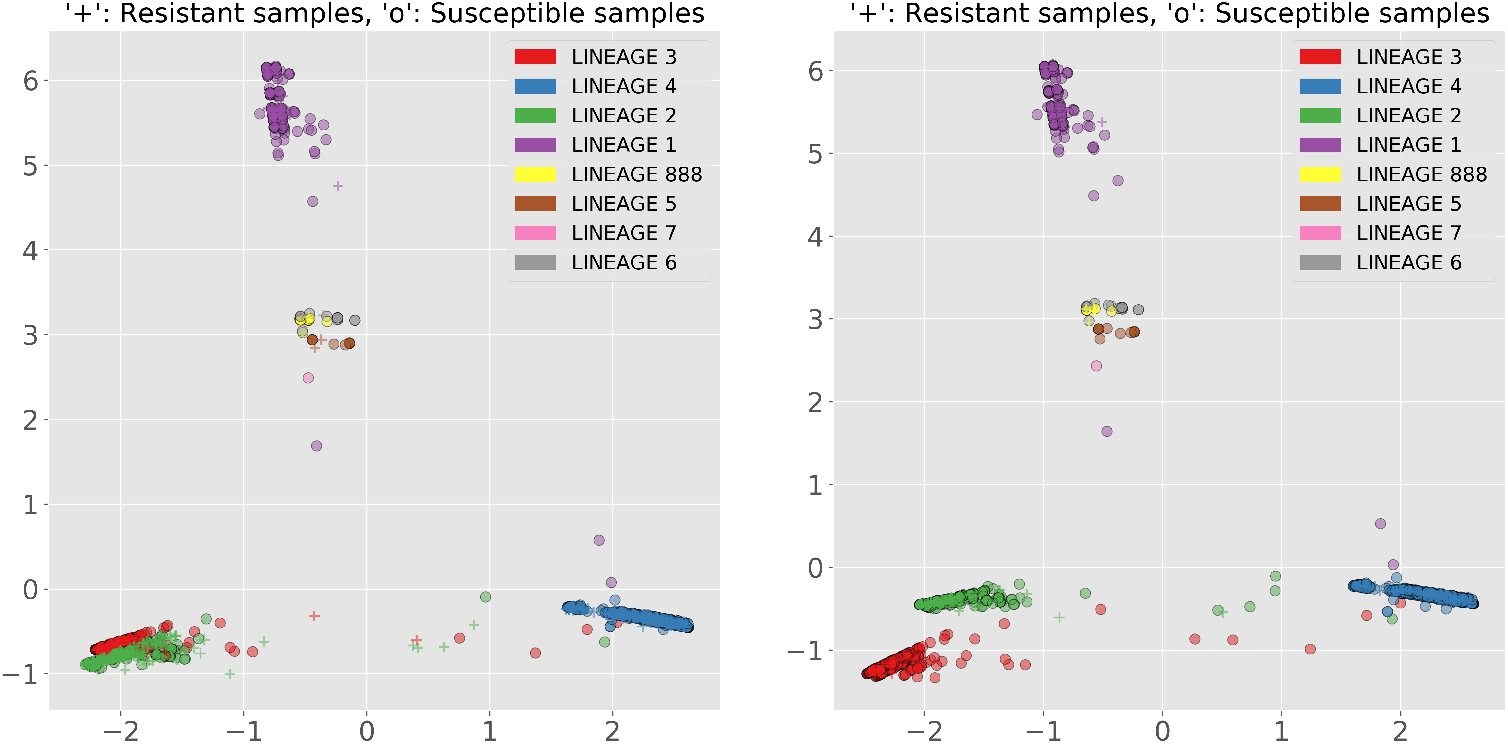
ReseqTB: The axes are the first 2 components of PCA. (Left) Isoniazid (Right) Pyrazinamide.

**Fig 6.**
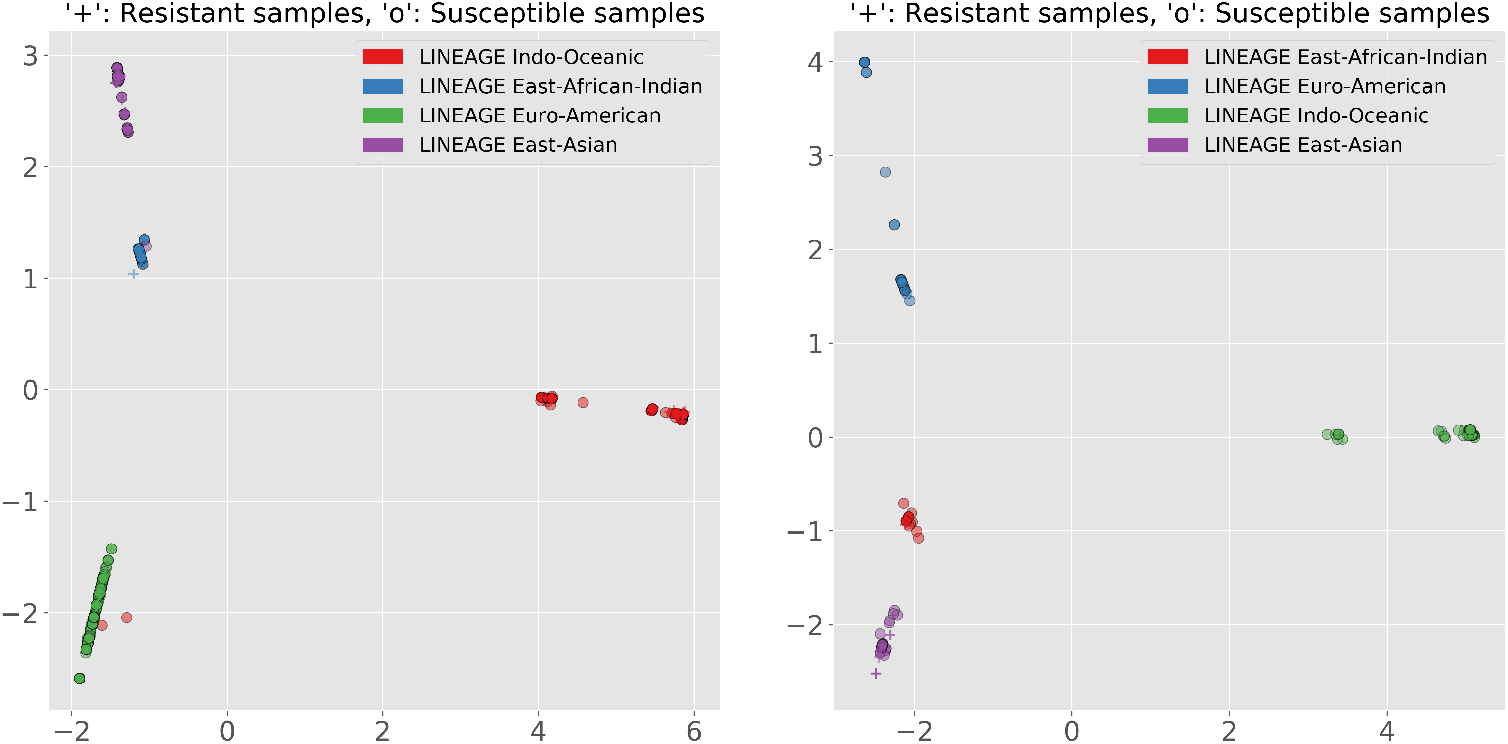
BCCDC:The axes are the first 2 components of PCA. (Left) Isoniazid (Right) Pyrazinamide.

### We achieved good performance when training and testing our models within a dataset

Any given probability threshold for assigning a prediction (drug-sensitive or drug-resistant) to each sample determines a sensitivity and a specificity for the model. Rather than using a fixed threshold, we take a holistic view and make use of the Receiver Operator Characteristic (ROC) curve. This curve can be used to determine the optimal sensitivity for a given specificity, or vice versa. Furthermore, one can use the Area Under this Curve (AUC) to evaluate and compare the performance of different models, as is commonly done in drug resistance studies [10, 20, 21].

#### Within-dataset performance differs between homogeneous and heterogeneous datasets

Test AUC plots are presented in Figure 7. In general, no model clearly outperforms the others. PZA is the most challenging drug to predict for the heterogeneous datasets PATRIC and ReSeqTB. For the latter, a larger dataset does not aid in PZA prediction. There appears to be a large variability in the mutations driving PZA resistance that makes it challenging for models to learn from a heterogeneous dataset, although PZA prediction performance is high for the homogeneous BCCDC dataset. This underwhelming performance may be partially explained by our genotyping method that does not take into account insertions and deletions, which can be determinant for PZA resistance [22].

**Fig 7.**
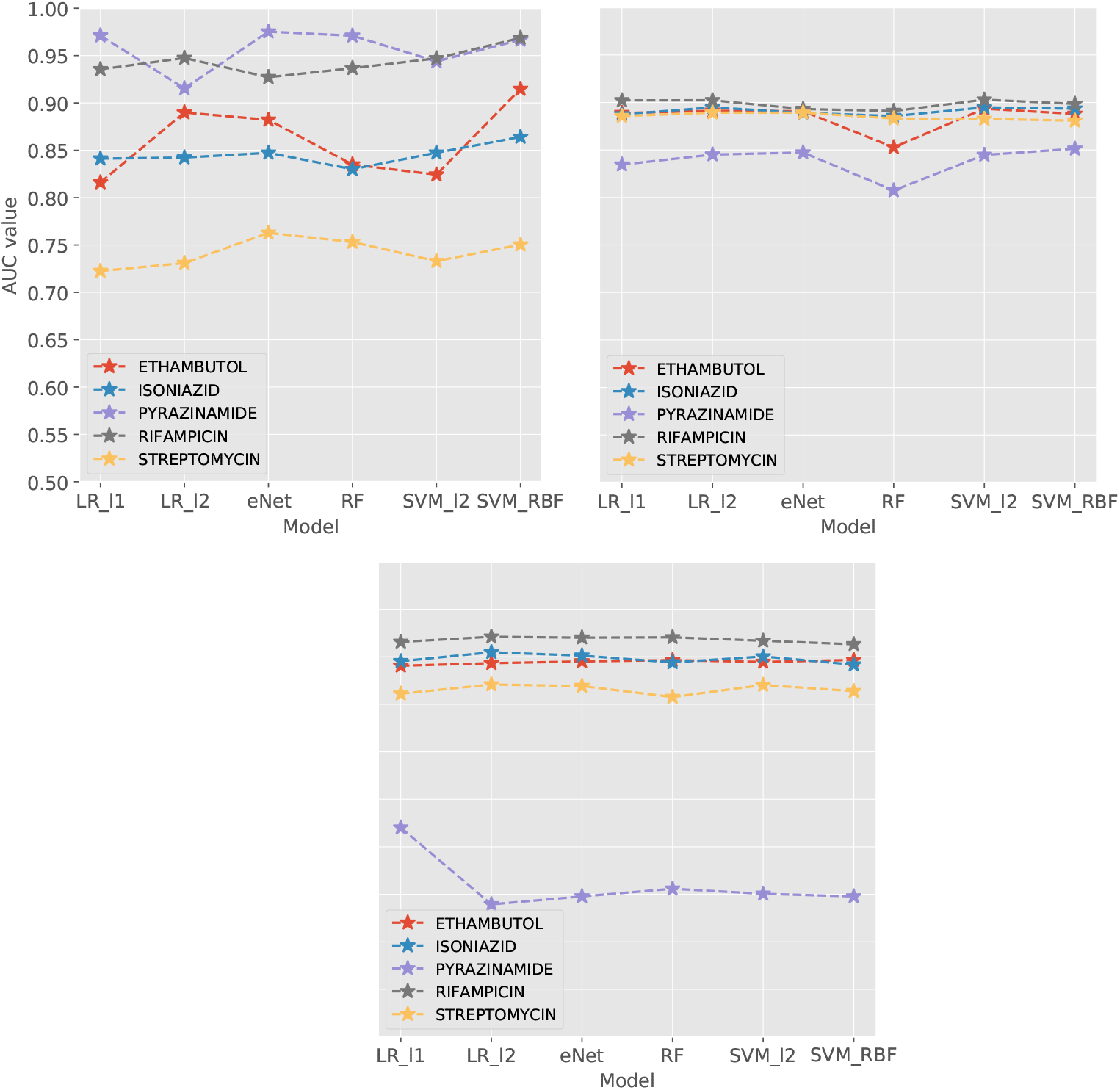
Test AUC for each dataset, trained on SNPs within 23 genes of interest. (Top left) BCCDC (Top right) ReSeqTB (Bottom) PATRIC.

Supplementary Figure 1 illustrates the results of training and testing within a dataset when training on the SNPs in 179 genes of interest. In general, no model clearly outperforms the others. PZA is the most challenging drug to predict for the heterogeneous datasets PATRIC and ReSeqTB. For the latter, a larger dataset does not aid in PZA prediction. There appears to be a large variability in the mutations driving PZA resistance that makes it challenging for models to learn from a heterogeneous dataset, although PZA prediction performance is high for the homogeneous dataset BCCDC. Moreover, the poor performance can be explained by the our method which does not take into account insertions and deletions, which can be determinant for PZA resistance [22].

The performance ranking of the heterogeneous datasets is identical, whereas BCCDC has PZA and RIF as the easiest drugs to predict, and SM as the worst one. Moreover, RIF and PZA are the easiest to predict for BCCDC, with AUC’s greater than 0.95, while SM is harder to predict than it is for any other dataset. The results obtained when training the models on SNPs within 179 genes are very similar to those on SNPs within 23 genes, with no obvious change in performance between the two. These results are shown in the Appendix.

### Cross-dataset evaluation shows the pitfalls of a model trained on a homogeneous dataset

By training our models on one dataset and testing them on another, we were able to observe how generalizable our models are. We use the short-hand notation *a → b* to denote a model with training set *a* and testing set *b*. For most drugs, prediction performance is higher when training on a heterogeneous dataset (i.e. ReSeqTB or PATRIC) and testing on a homogeneous dataset (i.e. BCCDC) than vice versa. We call training on a homogeneous dataset task A and training on a heterogeneous dataset task B. The results are shown in Supplementary Figure 2.

The gap in performance between task A and task B is larger for PZA than for the other drugs. There is an increase of 11% in the AUC of PZA while there is an increase of only 2% for RIF and even a decrease of 8% for ReSeqTB→BCCDC compared to its opposite. Nearly identical results are obtained for PATRIC→BCCDC and BCCDC→PATRIC. It is interesting to note that for PZA prediction, training on a heterogeneous dataset is better, while for SM and INH, training on a homogeneous dataset is better. Therefore, it is important to keep in mind not only the heterogeneity of a training dataset, but also the target drug.

We also found that training with 179 genes instead of 23 can in some cases make the model perform better in cross-dataset evaluation. For example, for ReSeqTB→BCCDC, the AUC for RIF increased from 73% to 83% when training on 179 genes. However, it is also possible for the performance to become worse, which can be seen for SM and INH.

### Cross-regional evaluation shows the pitfalls of applying a model trained on regional data on a different region

We also hypothesized that regional variation can have an effect on the predictive power of models, and thus in this section, we further investigate the regional impact of a model’s predictive power. The results are presented in Supplementary Figure 4.

Similar to our within-dataset evaluation, simpler models such as logistic regression outperform more complex models in most cases. For drug-specific performance, RIF is the most stable drug, meaning that the model can predict its resistance with relative ease throughout different regions. INH closely follows RIF, while PZA prediction fluctuates the most between regions. For a given model, PZA prediction performance can range from 50% to 90%, while RIF prediction performance is stable around 90%. It may be possible that mutations encoding for RIF and INH resistance are homogeneous across strains, meaning there is not much variation in the resistance mechanism across strains, while regional variation may exist for the resistance mechanism of other drugs such as PZA or SM.

Trends were also observed when training on different regional subsets. The North American model was much better at predicting resistance for African samples than Asian samples. Similarly, the African model was much better at predicting resistance on North American than Asian samples. The Asian model performed better on the North American than African samples, although the training data was limited. It appears that prediction of Asian samples may be a harder task from both the North American and African samples. It is possible that North American and African strains are genotypically closer to each other than Asian strains. However, this hypothesis would require further investigation and a much more comprehensive dataset, as each regional subset may not be representative of real-world strains present in each region. Furthermore, we did not use European data because the European subset appears to contain many different types of MTB due to the presence of many different groups of immigrants.

### Which genes are associated with drug resistance?

#### Submatrix permutation tests show that including more features may dilute the importance of key features

We considered different machine learning models for estimating gene importance using random permutation of the rows of the feature matrix, to provide an unbiased perspective. We chose the LR_L1, LR_L2, SVM_L2, SVM_RBF and RF models for this study and posited that a gene was important for resistance prediction if all the models assigned it a relatively high importance. We define *Delta* as the largest positive difference in test AUC between all models before and after permuting. The results are shown in Supplementary Table 1. We consider genes that are predicted to be important through machine learning as well as associated with resistance for the corresponding drug in [23] to be resistance-determinant.

For all three datasets, including 179 genes over 23 genes generally results in two scenarios: a decrease in the number of resistance-determinant genes deemed as important for resistance prediction, and/or a decrease in the *Delta* for resistance-determinant genes. For BCCDC (Table 2) with 23 genes, EMB, INH, RIF and SM the gene with the largest *Delta* (*Delta* > 10%) is a resistance-determinant gene. This trend remains when including 179 genes, however *Delta* < 10% for each case.

**Table 1.** Summary of datasets.

|  | Sample size | Countries | Number of SNPs<br>in 23 genes | Number of SNPs<br>in 179 genes |
| --- | --- | --- | --- | --- |
| **ReSeqTB** | 3428 | 28 | 2065 | 7351 |
| **PATRIC** | 1440 | 19 | 2793 | 8968 |
| **BCCDC** | 1138 | 1 | 734 | 2694 |
| **Overall** | 6006 | 39 | 4808 | 16 419 |

**Table 2.**
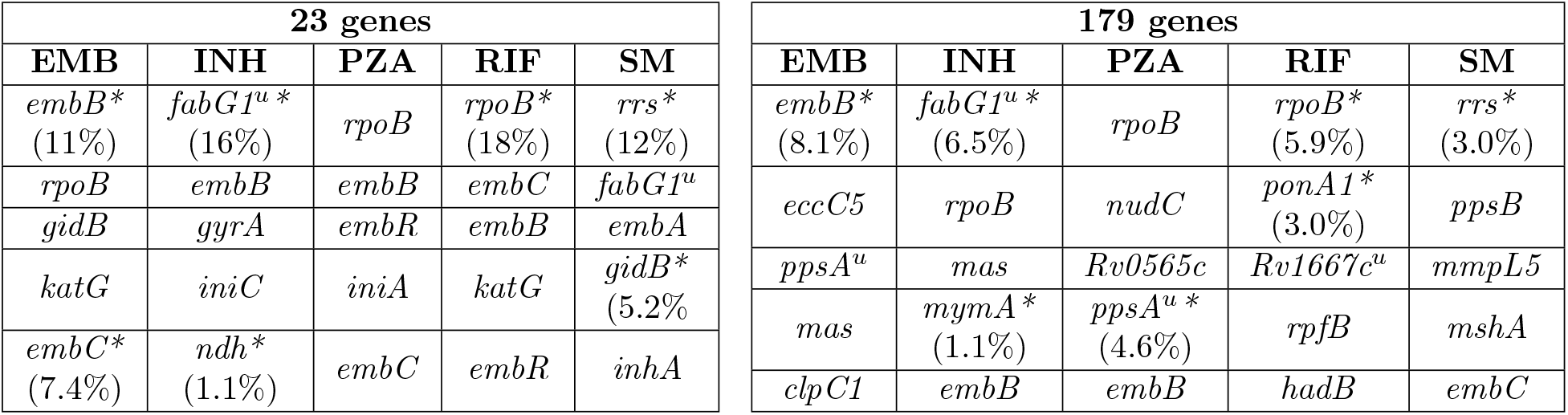
Top 5 genes with the highest *Delta* in the submatrix permutation tests in BCCDC. ^*u*^ indicates upstream regions and * indicates a gene known to be resistance determinant.

**Table 3.**
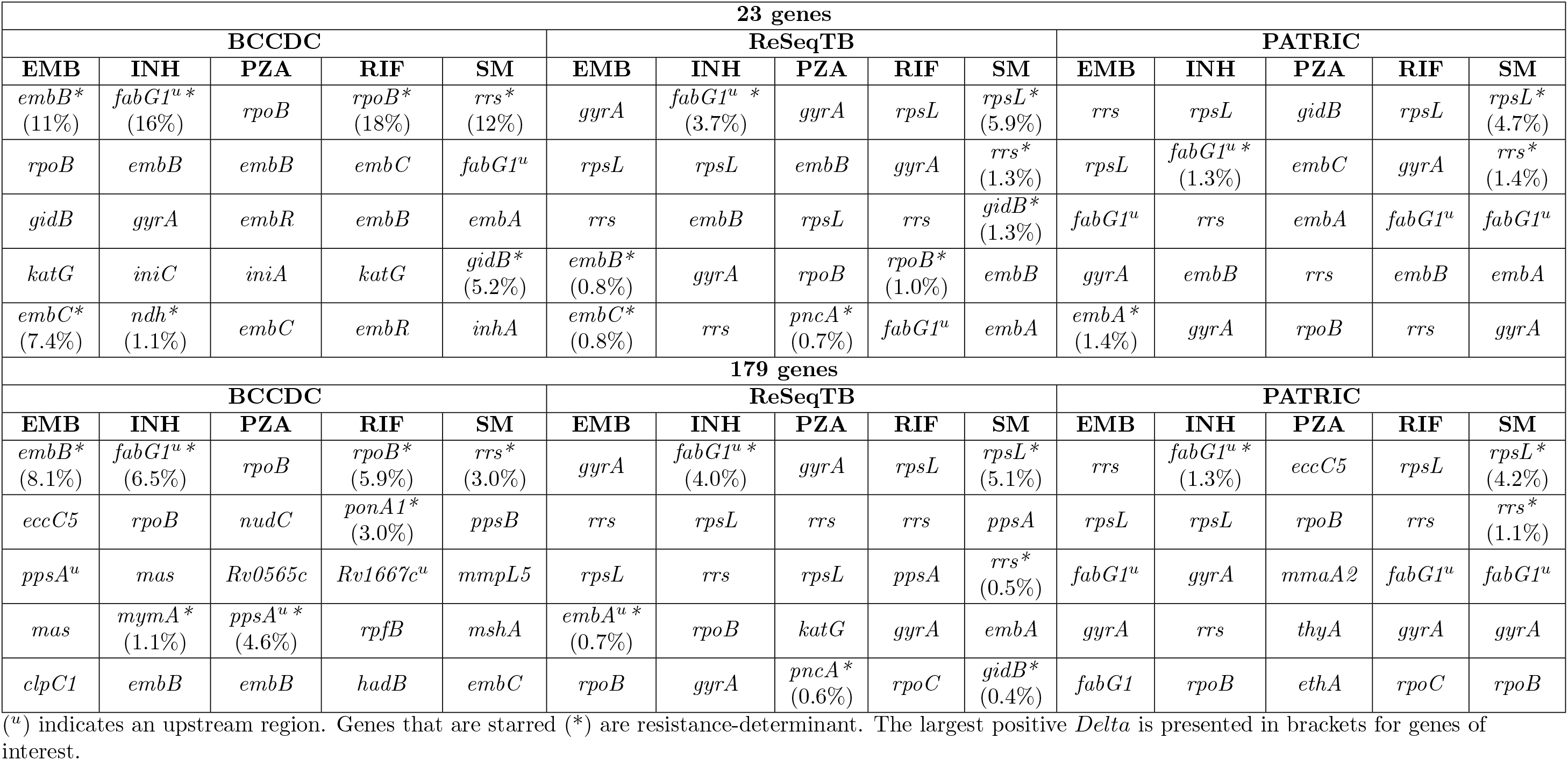
Genes with the highest *Delta* in the submatrix permutation tests.

**Table 4.**
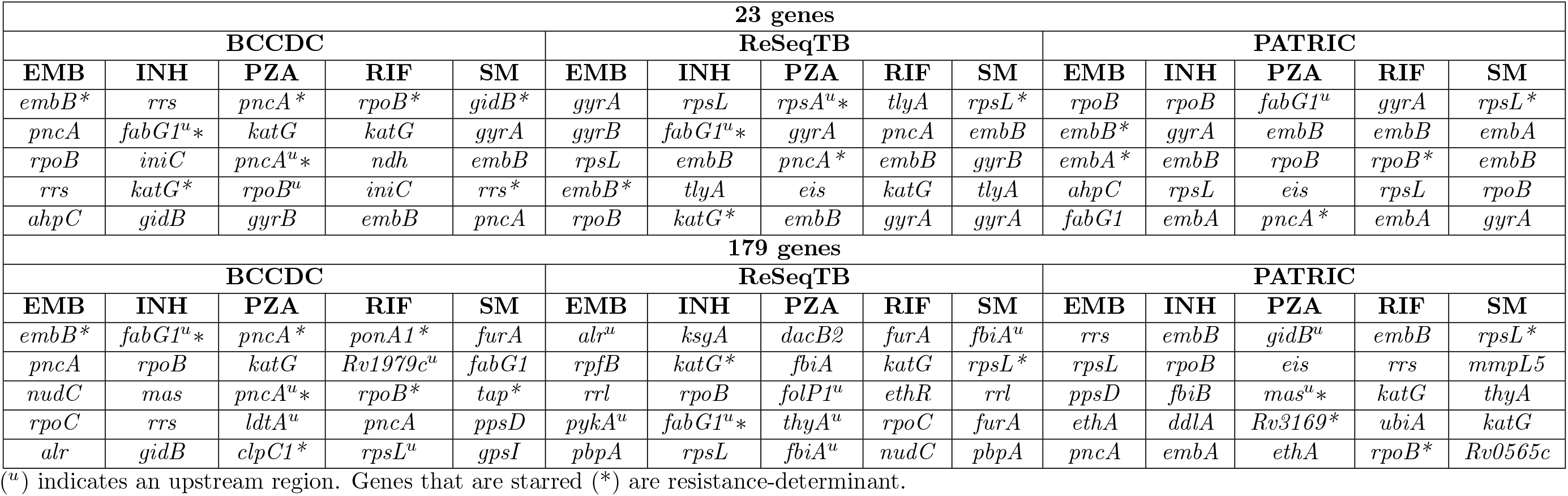
Genes with the largest coefficients in the LASSO model

**Table 5.**
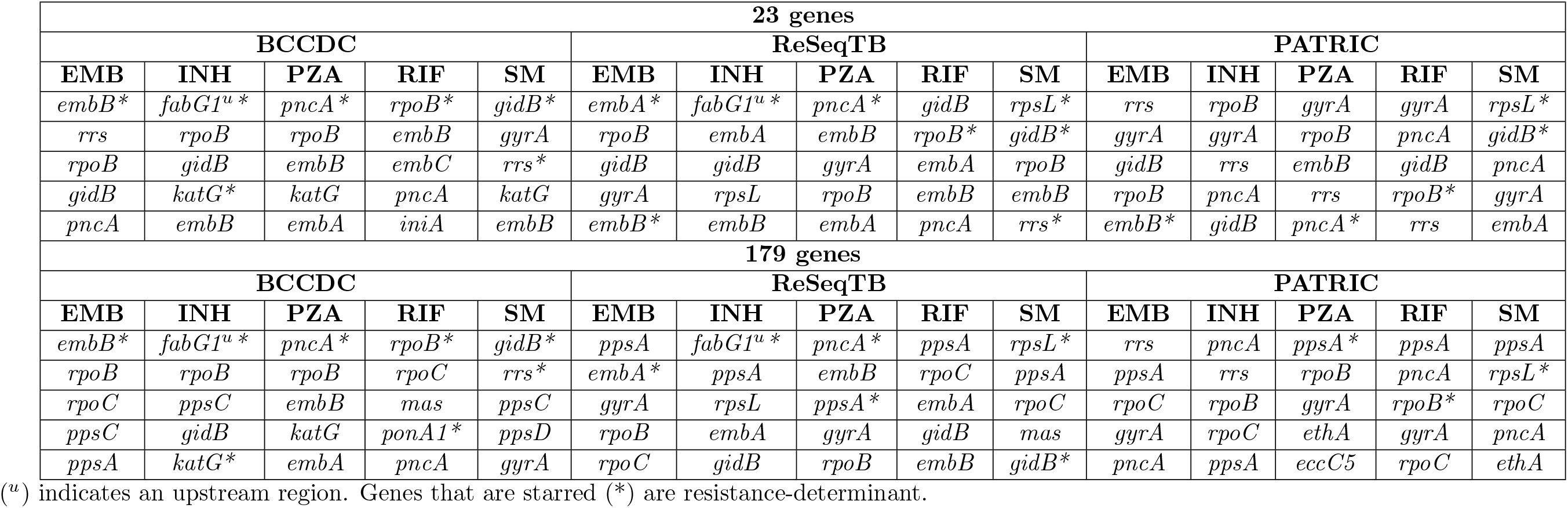
Genes with the largest importance using Random Forests

Moreover, we see that resistance-determinant genes are more easily captured as important in a homogeneous dataset. BCCDC has 4 drugs which have a resistance-determinant gene with the largest *Delta*, whereas ReSeqTB and PATRIC, the more heterogeneous datasets, only have this for at most two of the drugs each. The more modest performance for heterogeneous datasets may be due to a larger pool of possible variants. This hypothesis is supported by looking at the number of genes that have the five largest *Delta* values for each drug in PATRIC, the most heterogeneous dataset; there is a lower number of resistance-determinant genes identified among the top 5 than for any other dataset, perhaps due to the large number of different SNPs present.

#### Gene-wise partial least squares using the LASSO model reveals the cause of lower performance in heterogeneous datasets

We reduce the feature matrix gene-by-gene using PLS and end up with a transformed matrix with 23 or 179 columns, each representing a gene, which is then used as the input to a LASSO classifier. The non-zero coefficients of the LASSO and their magnitude can be interpreted as an estimate of the corresponding genes’ importance for resistance. We picked the regularization type by looking at the best performing logistic regression model for within-dataset evaluation. The results are shown in Supplementary Table 2.

A resistance-determinant gene is either the most or second most important feature for every drug in BCCDC. As the datasets get more heterogeneous, the models are less able to capture resistance-determinant genes as important. Only SM has a resistance-determinant gene as the most important for PATRIC, the most heterogeneous dataset. Moreover, including more features dilutes the importance of determinant genes. Many resistance-determinant genes identified within 23 genes are lost completely or have a lower importance when looking at 179 genes. There are cases where new resistance-determinant genes are found when including more features, but at the cost of reducing the importance of resistance-determinant genes.

#### Gene-wise partial least squares using Random Forests shows the difficulty of working with limited sample size heterogeneous datasets

We used the same reduced matrices as for the LASSO model, but as an input for RF. A gene’s importance was evaluated by investigating the model’s feature importance values. The results are presented in Supplementary Table 3.

The difference between using 23 and 179 genes is not as evident, but there is still evidence of resistance-determinant genes being masked when more features are introduced. The difference between a homogeneous dataset and heterogeneous dataset however, is clearer. BCCDC has a resistance-determinant gene as the most important gene for every drug. ReSeqTB has four drugs with resistance-determinant genes identified as the most important when including 23 genes, and three when including 179 genes. PATRIC, the most heterogeneous and the smallest dataset, only has one drug with a resistance-determinant gene as the most important for both 23 or 179 genes. This demonstrates the challenge of obtaining interpretable results with a small heterogeneous dataset.

## Discussion

In summary, we predicted TB drug resistance using different machine learning models on 3 large datasets, ReSeqTB, PATRIC and BCCDC, and performed the study of resistance-associated genes in a machine learning context. We also performed a cross-dataset evaluation and cross-regional evaluation.

The first contribution of our work is demonstrating the robustness of models when training and testing on datasets of different diversity. The test AUC for BCCDC is much higher than the ones for ReSeqTB or PATRIC. This demonstrates the ease of training and testing on a homogeneous dataset compared to a heterogeneous one. Furthermore, the performance difference between training on SNPs within 23 genes and SNPs within 179 genes was not significant, so increasing the size of the feature set does not consistently result in better performance when the sample size remains constant. Increasing the number of features may introduce too much noise that results in the model not performing any better. We also see that a model trained on a homogeneous dataset may not generalize well to a heterogeneous dataset. We demonstrate the potential difficulty of achieving good prediction results when a model trained on a homogeneous dataset is used to make predictions on a heterogeneous one. It is also important to keep in mind that the prediction of different drugs benefit from different level of heterogeneity in a dataset. We also demonstrate the potential challenges of training a model using data from one region. Such a model may only be suitable for that region, but may not be generalizable to other regions.

Apart from achieving good results on prediction, it is equally important to study the trained models and understand resistance-associated markers, something that seems to be a limitation of most previous work. Our second contribution is a rigorous method for identifying putative resistance-associated genes by permuting the rows of the submatrix of the corresponding gene as well as performing gene-wise PLS with LASSO. The gene-wise PLS method tends to give a better view of which genes are important, while the submatrix permutation testing method tends to be noisier and less consistent. However, our analysis still showed that some of the highly ranked genes were previously reported as resistance-determinant, supporting the use of our methods. Some other identified genes have not been extensively studied in conjunction with drug resistance, and thus may contain true signal or just noise. Therefore, one key challenge is to construct a reliable feature importance extraction method, preferably from a machine learning perspective, but possibly involving expertise-based feature engineering, to eliminate SNPs that may contribute to irrelevant genes being identified as important. By effectively eliminating noise, we are also able to include more relevant SNPs, which will potentially amplify the advantage of training models on a diverse dataset.

Our work can be seen as an exploratory study. To the best of our knowledge, it is the first one to investigate the issues of dataset homogeneity and heterogeneity and the identification of drug resistance markers at different resolutions by analyzing machine learning models trained on 3 major datasets and two sets of genes. We hope that this study will inform future work on drug resistance in pathogenic bacteria and the application of machine learning to the drug resistance problem.

## Supporting information

**List of 23 genes:** ahpC, fabG1, inhA, katG, ndh, rpoB, embA, embB, embC, embR, iniA, iniC, manB, rmlD, pncA, rpsA, gyrA, gyrB, rpsL, gidB, rrs, tlyA, eis

**List of 179 genes:** gyrB upstream, **gyrB**, gyrA upstream, **gyrA**, pbpA, ponA1 upstream, ponA1, ldtA, ldtA upstream, Rv0191 upstream, Rv0191, ldtE upstream, ldtE, inbR, inbR upstream, **iniA**, **iniC**, fgd1 upstream, fgd1, lprQ upstream, lprQ, mshA upstream, mshA, Rv0565c, Rv0565c upstream, hadA upstream, hadA, hadB, hadC, mmaA4, mmaA4 upstream, mmaA3, mmaA3 upstream, mmaA2, mmaA2 upstream, rpoB upstream, **rpoB**, rpoC upstream, rpoC, mmpL5, mmpS5, mmpR upstream, mmpR, rpsL upstream, **rpsL**, rplC upstream, rplC, rpfB upstream, rpfB, ksgA, mshB upstream, mshB, fbiC upstream, fbiC, sigI upstream, sigI, folP2 upstream, folP2, tap, tap upstream, **embR**, embR upstream, atpE upstream, atpE, atpC upstream, atpC, rrs upstream, **rrs**, rrl upstream, rrl, ldtD upstream, ldtD, fabG1 upstream, **fabG1**, inhA upstream, **inhA**, pykA upstream, pykA, rpsA upstream, **rpsA**, Rv1667c, Rv1667c upstream, **tlyA**, cycA, cycA upstream, eccB5 upstream, eccB5, eccC5, **ndh**, ndh upstream, **katG**, katG upstream, furA, furA upstream, Rv1979c, Rv1979c upstream, erm(37) upstream, erm(37), **pncA**, pncA upstream, blaC, blaC upstream, mshC, mshC upstream, kasA, eis, eis upstream, crfA upstream, crfA, ahpC upstream, **ahpC**, folC, ldtB, ldtB upstream, pepQ, pepQ upstream, Rv2671 upstream, Rv2671, Rv2731, thyX, thyX upstream, dfrA, dfrA upstream, thyA, thyA upstream, ald upstream, ald, gpsI, gpsI upstream, dacB2 upstream, dacB2, ppsA upstream, ppsA, ppsB, ppsC, ppsD, mas, mas upstream, ddlA, ddlA upstream, Rv3008 upstream, Rv3008, serB2, serB2 upstream, mymA upstream, mymA, Rv3169, whiB7, whiB7 upstream, nudC, nudC upstream, fbiA upstream, fbiA, fbiB, **manB**, **rmlD**, alr, alr upstream, rpoA, rpoA upstream, ddn upstream, ddn, nat, nat upstream, clpC1, clpC1 upstream, panD, folP1, folP1 upstream, proZ, proZ upstream, aftA, **embC**, embA upstream, **embA**, **embB**, ubiA, ubiA upstream, ethA, ethA upstream, ethR, **gidB**, gidB upstream

**S1 Fig.**
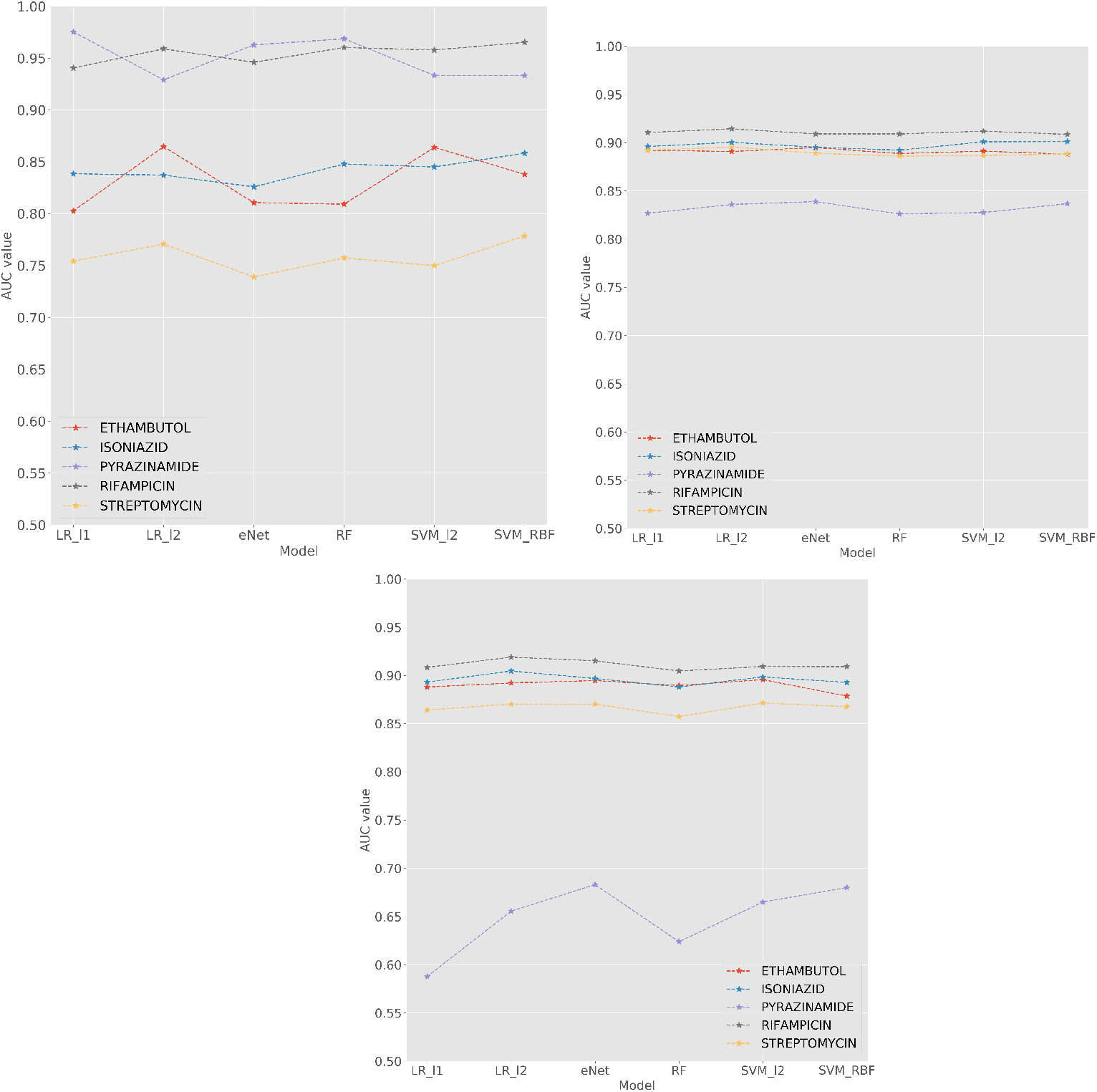
Test AUC for each dataset, trained on SNPs within 179 genes of interest. (Top left) BCCDC (Top right) ReSeqTB (Bottom) PATRIC.

**S2 Fig.**
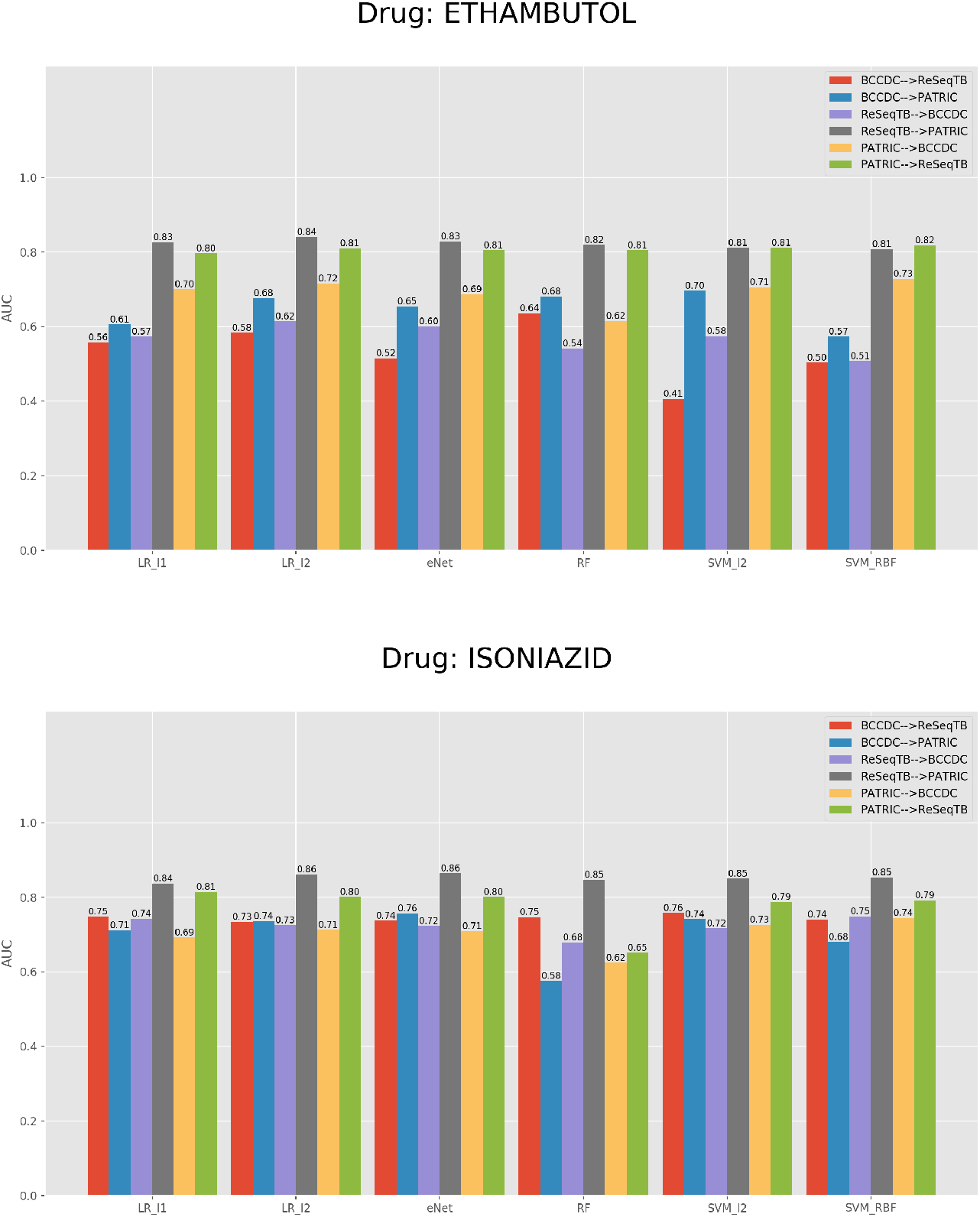
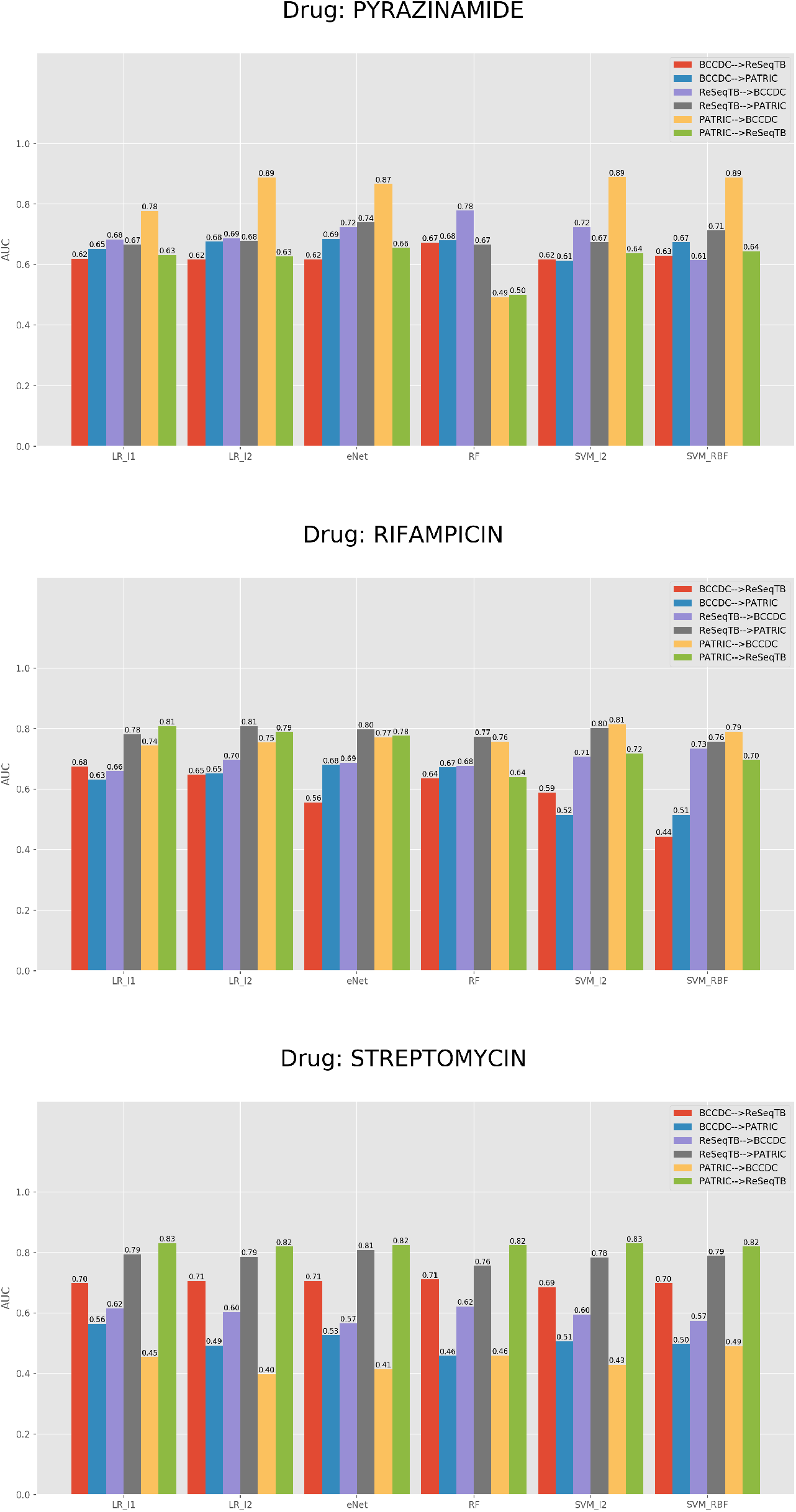
Test AUC for cross-dataset evaluation with SNPs within 23 genes of interest.

**S3 Fig.**
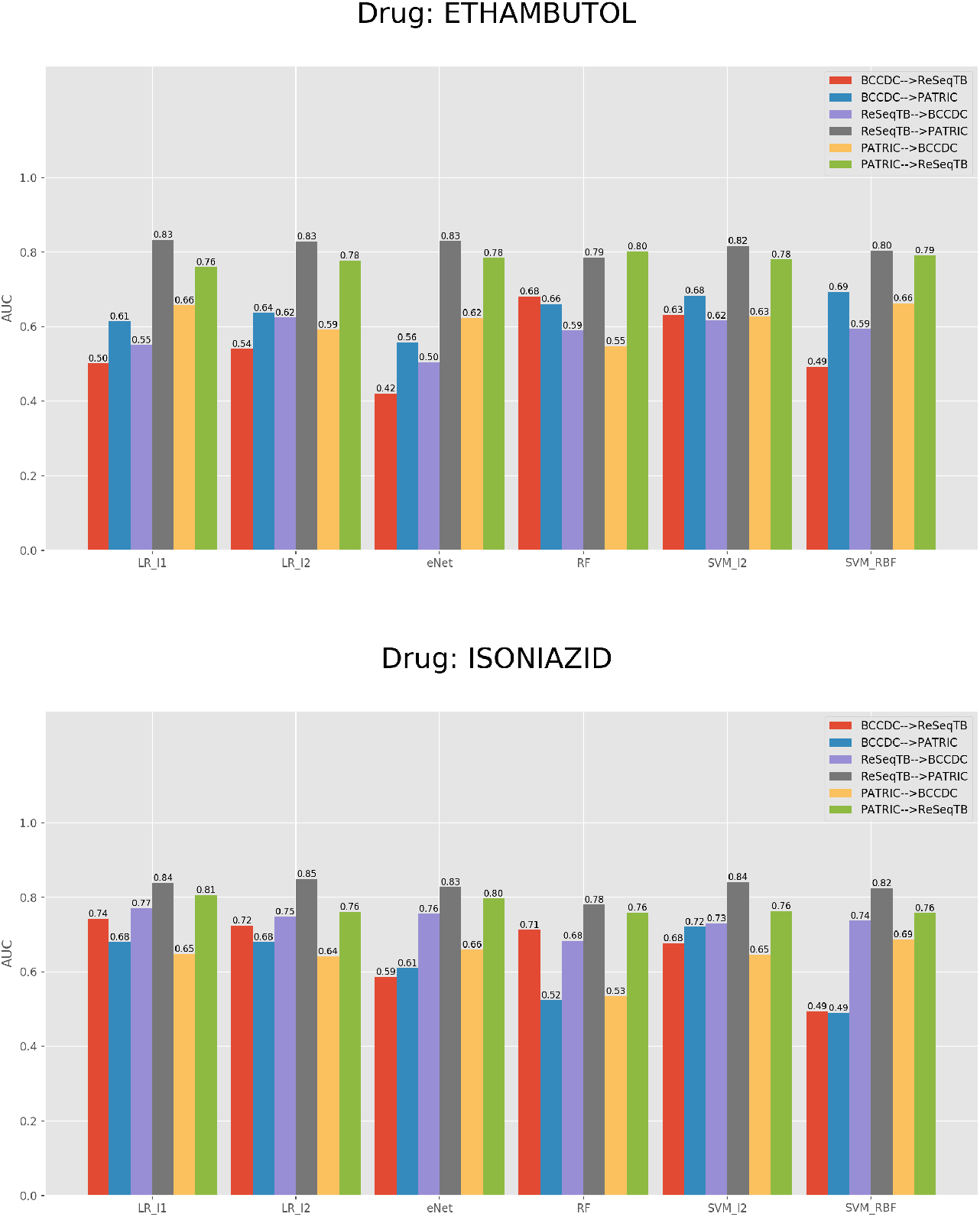
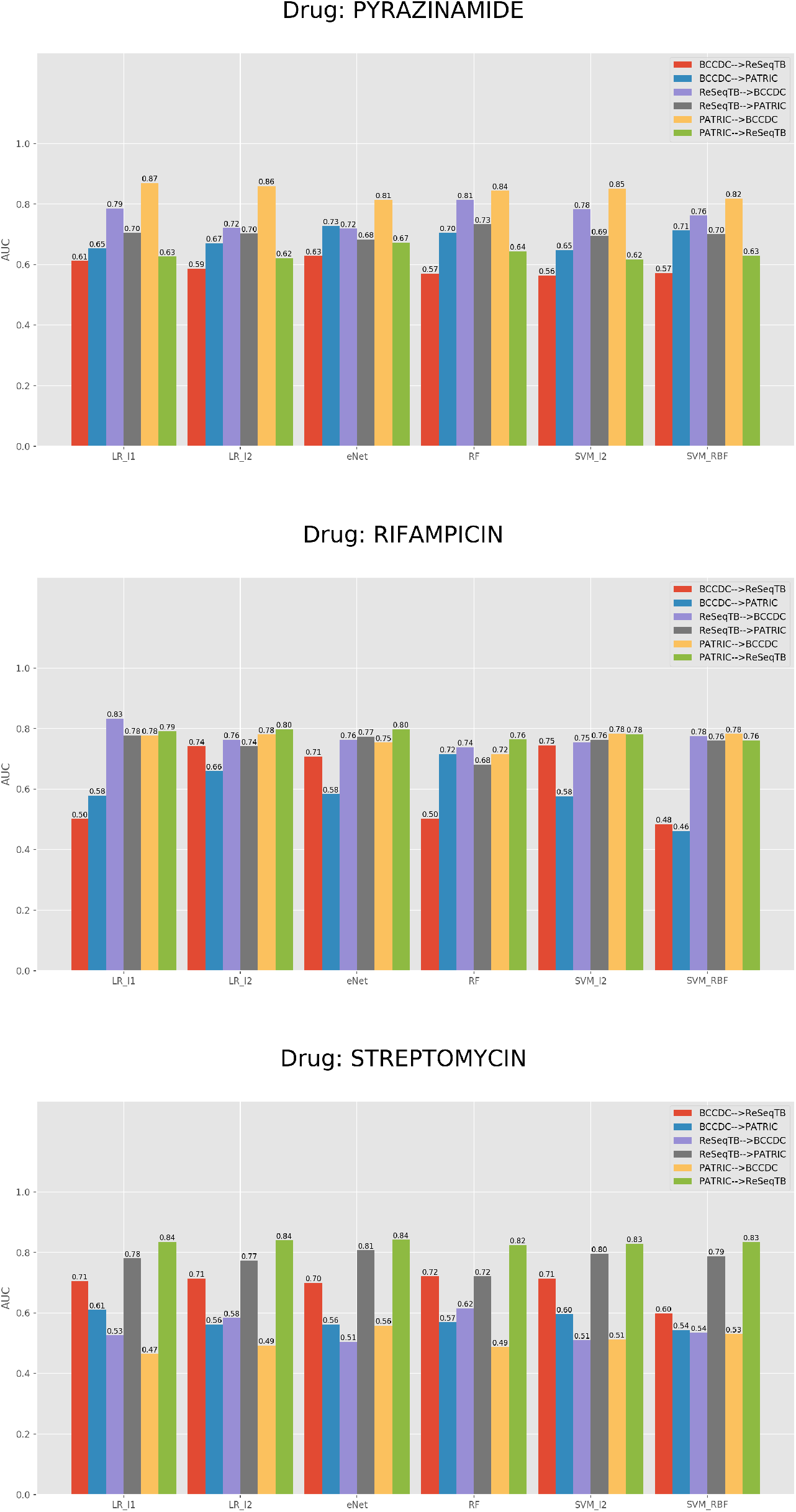
Test AUC for cross-dataset evaluation with SNPs within 179 genes of interest.

**S4 Fig.**
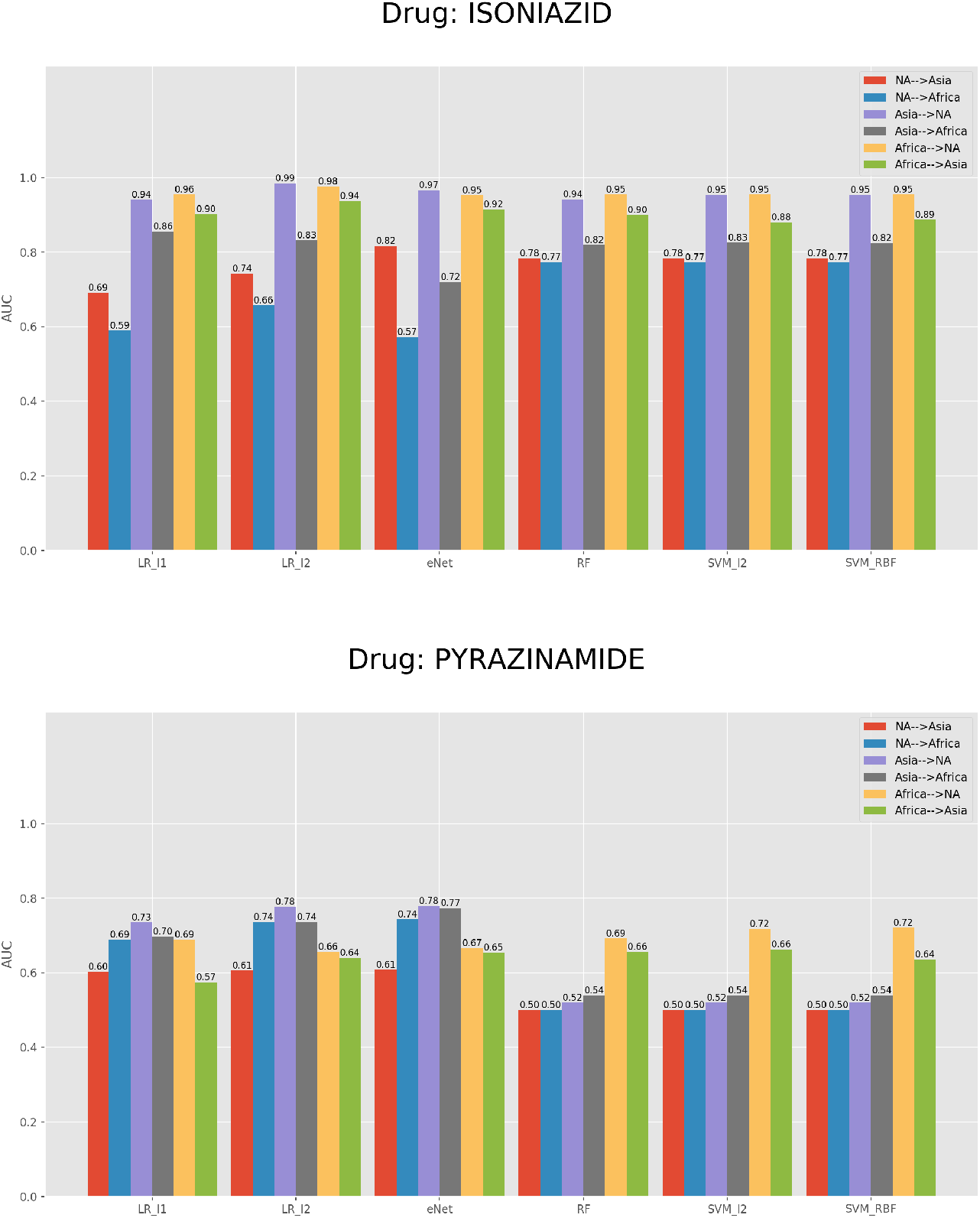
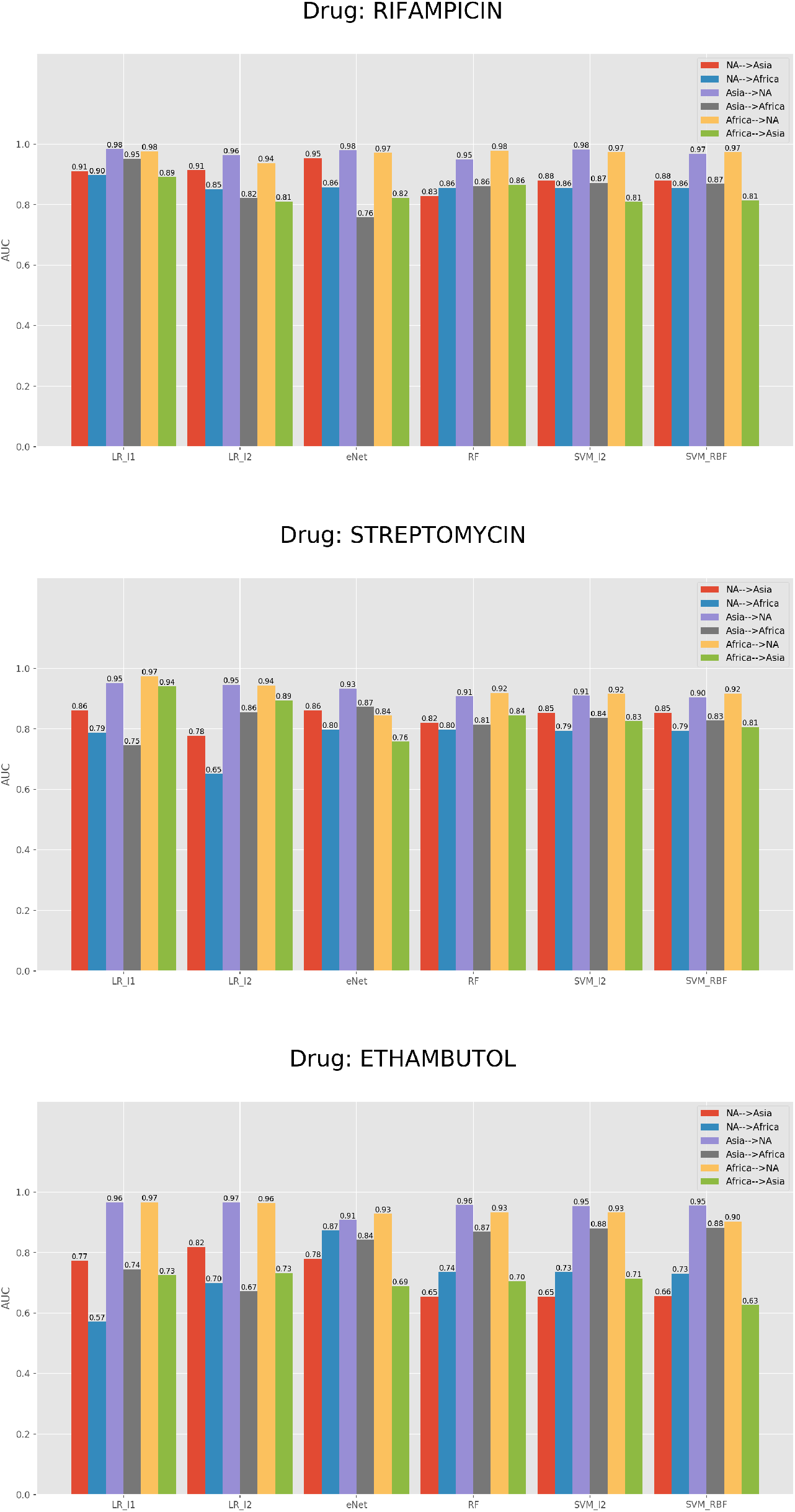
Test AUC for cross-regional evaluation.

**S1 Table. Top 5 genes with the highest *Delta* in the submatrix permutations tests for each dataset.**

**S2 Table. Top 5 genes with the highest LASSO coefficients for each dataset.**

**S3 Table. Top 5 genes with the highest random forests importance values for each dataset.**

## Acknowledgments

We would like to acknowledge Genome Canada, Genome BC and Genome Quebec for their support of Project code 283BAC that this paper is part of. We would also like to thank the BCCDC for help accessing their data and Canada’s Michael Smith Genome Sciences Centre for access to their compute clusters.

## Footnotes

1 Strains from the same lineage are generally thought to have originated from the same geographical location; they tend to share some unique mutations.

## Notes

### Competing Interest Statement

The authors have declared no competing interest.

### Summary of Updates

- This version addresses some concerns from a first round of reviews received for this paper

